# Synthesis of alkyl glucosides catalyzed by immobilized α-amylase from *Thermotoga maritima*

**DOI:** 10.1101/2024.06.28.601278

**Authors:** Wendy Xolalpa-Villanueva, Fidel O. Ramírez-Amador, Leticia Olvera, Alfonso Miranda-Molina, Agustín López-Munguía, Gloria Saab-Rincón

**Affiliations:** Departamento de Ingeniería Celular y Biocatálisis, Instituto de Biotecnología, Universidad Nacional Autónoma de México. Avenida Universidad 2001, Colonia Chamilpa C.P. 62210. Cuernavaca, Morelos, México; SYNMIKRO Research Center and Department of Chemistry, Philipps-University of Marburg, Marburg, Germany, Karl-von-Frish-Strasse 14, 35032, Marburg, Germany

**Keywords:** thermostable amylase, alcoholysis, alkyl glucoside, enzyme immobilization

## Abstract

The enzymatic production of alkyl glucosides is limited by the stability of the enzymes in the presence of alcohols. In the present study, we investigated three different supports: Sepharose 4B, crosslinked Sepharose (Fast Flow), and Eupergit C for the immobilization of α-amylase from *Thermotoga maritima*, AmyA, to increase its stability during the alcoholysis reaction. The enzyme immobilized on crosslinked Sepharose showed the best results, allowing its reutilization for at least five cycles while maintaining more than 50% residual activity. In addition, the immobilization of a previously reported H222Q variant resulted in a superior transglycosidic activity (> 50%) when compared to that of the wild-type (WT) AmyA. Moreover, both versions of the enzyme increased their residual activity after 24 h of incubation at 85°C upon immobilization. The alcoholysis with *n*-butanol, *n*-hexanol, and *n*-octanol was investigated to optimize the reaction conditions. Here, the addition of DMSO had a positive effect on the alcoholysis reactions with AmyA WT, achieving a total octyl glucoside title 1.75-fold higher than that obtained in the absence of DMSO.

**Importance:** Alkyl glucosides are valuable, non-toxic surfactants primarily synthesized chemically. The development of biocatalysts for their production has become a significant goal in the field of biocatalysis to avoid the disposal of toxic waste and environmentally harmful processes. To make these processes competitive, the use of low-cost raw materials and the recycling of biocatalysts are essential. The immobilization of *Thermotoga maritima* α-amylase has enabled its use in the presence of long-chain alcohols, achieving octyl glucoside production of 0.7 mg/mL—an unprecedented feat for an amylase. This study represents a breakthrough in the use of α-amylases for alkyl glucoside production.

## Introduction

Alkyl glycosides or alkyl polyglycosides (APGs) are non-ionic surfactants synthesized by condensation of one or more carbohydrate units and a fatty alcohol; they are specifically named alkyl glucosides if the carbohydrate is a glucose molecule. Due to their chemical properties, APGs are suitable for applications in detergent, cosmetic, and pharmaceutical industries with almost negligible toxicity (1, 2). In formulated products, there is evidence of the significant effect of APGs as stabilizing surfactants of lipid nanoparticles in comparison to classical stabilizers, i.e., they are less prone to oxidation at room temperature than other surfactants such as polysorbates (3). The application of APGs in other fields is constantly growing. In the oil industry, APGs have been shown to enhance product recovery as a valuable chemical flooding surfactant (4, 5).

On top of their usefulness as industrial surfactants, APGs are biodegradable. Thereby, APGs are currently considered a new generation of bio-based surfactants or green surfactants, since many of them are derived from natural sources (i.e., palm kernel oil and coconut oil, among others). An increasing demand for these APGs has inspired the development of methods for their chemical or enzymatic synthesis. The traditional chemical methods for their synthesis are based on Fischer glycosylation, through the alcoholysis of glucose under acidic conditions, or by trans-acetylation (6). These methods, however, require harsh conditions as well as toxic organic solvents that are not amenable to the environment (7).

Alternatively, an enzymatic reaction represents a more sustainable and environmentally responsible option for the synthesis of APGs, as enzymes are both selective and stereospecific, they can use renewable feedstock as substrates, require fewer steps, and reduce the use of solvents and waste products (6). Leloir glycosyltransferases are the natural enzymes to carry out this reaction. Nevertheless, their need for activated sugars as donor groups makes them unsuitable for industrial purposes. For this reason, enzymes within the glycosyl hydrolase family have been investigated as potential transferase enzymes. Among the enzymes used to produce APGs are some glycosidases and members of the non-Leloir glycosyltransferase groups. In this regard, a β-galactosidase was reported to produce the equivalent of 66 mM butyl galactoside using lactose as a glycosyl donor substrate in a butanol system saturated with water (8). Importantly, both bacterial and fungal β-glycosidases, along with almond glycosidase, are commonly employed for APGs production by transglycosylation (9). An alternative glycosyl donor, like *p*-nitrophenol glucopyranoside (*p*NPG), has been used to facilitate the estimation of conversion percentage, reaching up to 97 % conversion with methanol by the Thai Rosewood β-Glucosidase (10); however, *p*NPG is an expensive substrate for industrial purposes and the residual *p*NP unacceptable. In contrast, α-amylases are an attractive alternative due to their ability to take starch – an abundant and inexpensive substrate – as a glycosyl donor. In particular, the α-amylase from *Aspergillus oryzae* has been tested with different alcohols as acceptors to produce methyl, ethyl, benzyl, propyl, and butyl glucosides (11). The transfer of glycosides by α-amylases is not restricted to monosaccharides as larger saccharides may also be transferred. Other products, such as methyl maltoside and methyl maltotrioside have also been detected when methanol is used as the acceptor. These by-products may be of interest per se to modulate the toxicity of the alkyl glycosides (12), but if the monosaccharide product is the goal, such by-products can be sequentially hydrolyzed to methyl glucoside, for instance by the amyloglucosidase from *Aspergillus niger,* as demonstrated by Larson and coworkers, who reported a 3.6 mM production of methyl glucoside, produced from the hydrolysis of the methyl maltoside and methyl maltotrioside main products (11). Although the use of amylases to produce APGs has undoubtedly been successful, the use of longer-chain alcohol acceptors, such as hexanol or octanol, would result in more robust surfactants; however, such glycosylations have been achieved mainly with β-glycosidases (9). Despite the potential of amylases, there are several challenges to overcome for their application as efficient biocatalysts. These include the operation at higher temperatures and alcohol concentrations required for a higher selectivity of the alcoholysis reaction. In this context, the discovery of new enzymes from thermophilic organisms represents an important source to explore the power of enzyme catalysis beyond their natural environment and common reactions. Amy A, a recombinant α-amylase from the hyperthermophilic bacterium *Thermotoga maritima*, is not only able to transfer glucose molecules from starch to methanol at 85 °C, but also tolerates alcohol concentrations up to 40 % (13). In this way, Amy A has been used to produce methyl and butyl glucosides reaching equilibrium concentrations of 31.7 mM and 16.9 mM, respectively (14). Additionally, the H222Q variant of AmyA has been reported to reach a two-fold increase in the alcoholytic efficiency compared to the WT enzyme (14). Nonetheless, optimization is required to scale up this process in a more affordable fashion.

Currently, well-standardized methods to produce recombinant enzymes are designed to obtain high protein yields at a relatively low cost, although reaching sufficiently pure samples for subsequent applications remains challenging. Therefore, reusing enzymes repeatedly during the process is advantageous and, in some cases, an absolute requirement. The immobilization of enzymes facilitates their reutilization for several catalytic cycles, provided an adequate immobilization support, and a suitable immobilization method are selected. Amylases from very different sources, especially those from bacteria, have been immobilized using a variety of supports such as alginate, agarose, chitosan, and other modern materials such as acrylic fibers (15) and magnetite alloys (16); however, in most cases the immobilized preparations have been tested only for hydrolysis reactions with no reports of immobilized amylases being used in the presence of alcohols for the synthesis of APGs. In this study, we investigated the alcoholytic activity of the immobilized α-amylase AmyA WT from *T. maritima* and its variant H222Q in butanol to produce butyl glucoside. Reactions with hexanol and octanol were also included in this study. Noticeably, the addition of co-solvents such as polyethylene glycol (PEG) or dimethyl sulfoxide (DMSO) showed a positive effect in improving the miscibility of long-chain alcohols on the alcoholytic reactions.

## Materials and methods

### Expression and purification of recombinant amylases

The cloning of the *amyA* gen from *Thermotoga maritima* and construction of the variant H222Q by site-directed mutagenesis was constructed as previously reported (14). Plasmids containing the WT and the mutated genes were transformed in the *E. coli* C41 (DE3) strain. The resulting colonies were picked from Luria Bertani (LB) agar plates with ampicillin 200 µg/mL and inoculated in LB liquid media with the same antibiotic. The expression of recombinant proteins was induced by the addition of 0.25 mM IPTG and incubation at 20 °C for 16 h. Cells were harvested and disrupted by sonication. The lysate was centrifuged for 30 min at 3800 X g at 4°C to separate the supernatant from the pellet, and the supernatant sample was further heated for 1 h at 70°C before being centrifuged to separate the soluble fraction. The protein sample was subjected to affinity chromatography on Ni-NTA agarose beads, and fractions containing the protein of interest were pooled and dialyzed against enzyme buffer (50 mM Tris, 150 mM NaCl, 2mM CaCl_2_, pH 7) and assessed for activity. For immobilization, the enzyme solution was dialyzed against 50 mM phosphate buffer and 150 mM NaCl at pH 7. The protein concentration was estimated by the bicinchoninic acid method (Pierce) and by Absorbance at 280 nm using the molar extinction coefficient ε 148170 M^-1^ cm^-1^.

### Assessment of hydrolytic activity

Hydrolysis reactions were performed in the presence of 1% starch and 15.6 nM of free purified protein WT or H222Q in a final volume of 1 mL at 85 °C and 500 rpm in a thermoblock mixer (Eppendorf). Soluble starch (Sigma) was gelatinized in enzyme buffer containing 50 mM Tris, 150 mM NaCl, and 2 mM CaCl_2_, pH 7 preheated at 85°C before starting the reaction. The reaction was followed over time by measurement of reducing sugars with the DNS method (17). Briefly, a 50 µL reaction sample was taken every minute and added immediately to an equal volume of DNS solution. The mixture was heated in a thermocycler at 95°C for 10 min and read the absorbance at 540 nm. The amount of reducing sugars was expressed as glucose equivalents by interpolating the absorbance data to a glucose standard curve. One activity unit (U) was defined as the amount of enzyme required to release 1 µmol of glucose per minute. Similarly, the specific enzyme activity was defined as the number of enzyme units per mg of protein (U/mg).

### Assessment of alcoholytic activity

Alcoholysis reactions were performed with 10% starch and 10 U of free enzyme in a final volume of 0.5 mL. To reach such a concentration, starch was prepared directly in a 2 mL tube and gelatinized at 85 °C in enzyme buffer containing 10% *n*-butanol, or in a mixture of the specified alcohol and either 5% PEG or 30% DMSO as co-solvents. All reactions were performed in a thermoblock mixer (Eppendorf) at 500 rpm and 85 °C for 24 h unless other conditions were specified. At the end of the reaction, a 0.5 µL were applied on a silica plate (silica gel 60 F_254_ Millipore), and the reaction products were separated by thin layer chromatography (TLC) using a mixture of ethanol:butanol:H_2_O (50:30:20 or 40:40:20) as the mobile phase. Both alcoholysis and hydrolysis products were revealed by spraying the dried plate with a solution of α-chloro naphthol and sulfuric acid before heating at 110°C with a heat air gun. The resulting spots were compared with glucose, butyl glucoside, or octyl glycoside standards. In the case of malto-oligosaccharides or alkyl oligosaccharides, these were converted to their monoglucosylated form by diluting 100 µl of the alcoholysis reaction with enzyme buffer (1:1) and digesting with 2.5 U of amyloglucosidase from *Aspergillus niger* (Sigma) at 40 °C for 6 h. TLC analysis was performed using one µl of the digested sample.

### Determination of alkyl glucoside concentration

The amount of alkyl glucoside present in the alcoholytic reactions was measured by high-performance liquid chromatography (HPLC) analysis of the amyloglucosidase digested sample. Samples were filtered through nylon 0.22 µm filters (Thermo) and injected with an automatic injector (Waters 717 Plus autosampler) onto a Phenomenex reverse-phase column. A Waters-Millipore liquid chromatography instrument equipped with a refraction index detector (Waters 410) was used. The mobile phase was set according to the mixture of products to be analyzed; methanol:H_2_O 10:90 phase was used for butyl glucoside, while methanol:H_2_O 60:40 phase was used for octyl glucoside. An α-octyl glucoside or butyl glucoside standard was used for comparison with the peak corresponding to an alkyl glucoside. Glucose, alcohols, and cosolvents were injected independently to analyze their retention times and to assess that they did not interfere with the quantification of the corresponding alkyl glucosides.

### Enzyme immobilization

Sepharose 4B CNBr activated (Cytiva), crosslinked Sepharose CNBr activated Fast Flow (FF, Cytiva), and Eupergit C (Röhm GmbH & Co. KG) resins were used as supports. Prior to immobilization, 30 mg of Sepharose, 50 mg of crosslinked Sepharose FF, or 50 mg of Eupergit C were prepared in a 2 mL tube. The supports were treated according to the manufacturer’s indications with a few modifications. Briefly, both Sepharose supports were washed three times filling the tube with cold 1 mM HCl, followed by a washing step with cold filtered water. Then, the samples were centrifuged at 3000 rpm for 2 min at room temperature in a benchtop microcentrifuge, before removing the supernatant. The washed beads were equilibrated with immobilization buffer consisting of 100 mM phosphate buffer and 500 mM NaCl, pH 7. The beads were immediately coated with a 1:1 volume solution of purified protein (1.0 or 2.4 mg/mL) in immobilization buffer and incubated for 24 h at room temperature in an orbital shaker. Unbound protein was removed by centrifugation and the protein concentration and residual activity were measured to estimate the amount of immobilized protein. The coupled sepharose was washed in 1 M ethanolamine for 2 hours at room temperature to block unused activated sites. To remove any non-covalently bound protein, the beads were washed with four alternating rounds of 100 mM Tris, 500 mM NaCl, pH 8, and 120 mM sodium acetate buffer, 500 mM NaCl, pH 3. Finally, the beads with immobilized protein were stored in an enzyme buffer at 4°C until use. For the Eupergit C matrix, the protein was diluted to 15.6 and 31.2 µM in 1 M potassium phosphate buffer pH 7.5, added directly to weighted Eupergit C matrix, and incubated at room temperature for 24 h on a rotating shaker. The solution with the unbound protein was isolated. Protein quantification and residual activity were determined. The Eupergit C beads with immobilized protein were washed 3 times with 1 M NaCl and equilibrated in enzyme buffer before storage at 4 °C. Three replicates of enzyme immobilization were performed on each matrix.

The immobilization yields were calculated as follows (18):

Enzyme activity yield (%) = 100 × (immobilized activity/starting activity)

The immobilized activity was obtained by subtraction of the remaining activity measured in the enzyme solution after immobilization from the starting activity.

The efficiency of immobilization is defined as the activity units of bound enzyme determined experimentally relative to the immobilized activity described above.

Efficiency of immobilization (%) = 100 X (Total activity in the immobilized protein/immobilized activity)

### Determination of Hydrolytic activity

For hydrolysis reactions, 5 µL of immobilized enzyme was applied to 1 mL of 1 % starch previously solubilized. The hydrolytic activity was followed as stated above for the free enzyme. The agitation in the thermoblock mixer (Eppendorf) was increased to 800 rpm, and 50 µL samples were taken at different time points to determine reducing sugars by the DNS method. Reactions were done by triplicate.

### Determination of Alcoholysis activity

For the alcoholysis reactions, 10 µL of immobilized enzyme was used in 10 % starch with 10% butanol or other alcohols and co-solvents as indicated before. The reaction was incubated at the specified temperature during 18 h, and alcoholysis products were determined as described above. After the first alcoholysis cycle, the soluble products were separated from the immobilized enzyme by centrifugation. The support was washed twice with enzyme buffer and the following cycle was started in a second tube containing fresh gelatinized starch. Alcoholysis reactions were done by duplicate.

### Temperature stability of the free and immobilized enzymes

Residual enzyme activity was measured by comparing the activity of the protein incubated at 65 °C or 85 °C, and room temperature (control) for 6 and 24 h. Here, either 0.1 mg/mL enzyme or 200 µL of beads containing the immobilized enzyme were used. After incubation, 10 µL enzyme at each condition were added to 1 mL of 1% solubilized starch (preincubated at 85°C and pH 7) to test residual activity. The production rate of reducing sugar was measured by the DNS method and the assays were performed in triplicate.

### pH profile of the free and immobilized enzymes

The pH activity profile of AmyA WT and H222Q was evaluated in the free and immobilized forms on the Sepharose FF support. The same conditions used to evaluate the free enzyme were used, but the pH was varied, using a mixture of buffer solutions with 100 mM phosphate-citrate-glycine containing 150 mM NaCl in a pH range from 5 to 10.5. The value corresponding to the maximum activity point observed with the free form was taken as 100 % of activity.

## Results

### Evaluation of the enzyme immobilization support

Since alcoholysis reactions require both high temperatures and the presence of alcohol, the first step consisted of the selection of a compatible support to enhance the enzymés performance under such conditions. We compared two commercially available matrices, Eupergit C and Sepharose. The former has been described as suitable for the covalent immobilization of enzymes for industrial applications (19), while the latter has been extensively used for chromatography (20), protein binding and pull-down-related applications (21). Table 1 shows the immobilization yields and efficiencies obtained with each support when three different amounts of AmyA WT were loaded. We tested two different forms of a CNBr-activated Sepharose matrix: non-crosslinked and crosslinked (also known as Fast Flow). The immobilization yield was obtained by subtracting the activity found in the flow-through and washings from the total activity loaded, corresponding to the non-retained enzyme. For all three supports tested, the immobilization yield was > 90 %. However, the immobilization efficiency of the Eupergit C — calculated from the activity directly measured in the support, relative to the amount of immobilized protein — was considerably lower in comparison to the Sepharose versions. The hydrolytic activity assays were evaluated using soluble starch solubilized at high temperature as a substrate. According to Liebl et al. (23), the optimal temperature of AmyA WT is around 85 °C and the optimal pH is 7. However, the enzyme immobilized in Sepharose 4B (non-crosslinked) did not withstand this temperature as agarose tends to melt above 80 °C. Importantly, when the temperature for the alcoholysis reactions was lowered to 60°C, we were able to successfully produce alkyl glucosides after 24 h, as shown by the TLC analysis, where the presence of butyl glucoside can be observed for several cycles (Figure S1).

**Table 1.**
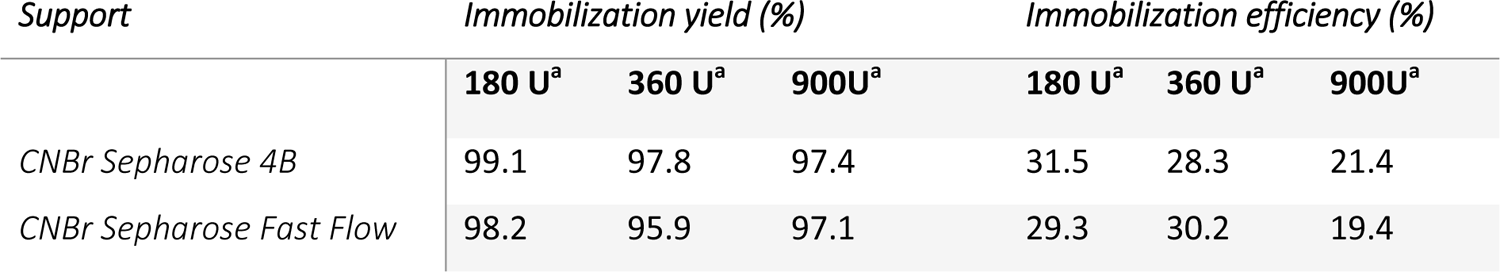
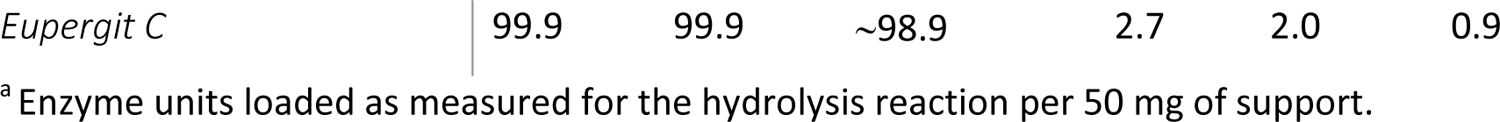
Evaluation of the immobilization process of AmyA WT on three different supports.

### Hydrolytic activity of immobilized Amy A

We investigated the immobilization support capacity by adding three different amounts of enzyme in the immobilization step. As a control, we tested the support treated under the same immobilization conditions but without any bound protein to ensure the absence of a background signal of any of the supports during the DNS assay (data not shown). Then the same amount of support with immobilized protein was employed in the hydrolysis reactions. Since the initial reaction rate was determined during a maximum of 10 min, we followed the evolution of the reaction during this time. As shown in Figure 1, the activity rate when 180 and 360 U of protein were loaded per 50mg of support increases accordingly. However, the immobilization efficiency — estimated by the experimentally determined activity vs the immobilized activity (18) — decreased at high loading suggesting that the protein active site was less accessible at this high protein concentration. On the other hand, a very low activity of 0.23 µmol/mL min^-1^ ± 0.06 is detected with the enzyme in Eupergit C, which did not show the same increase upon increasing the amount of loaded enzyme.

**Figure 1.**
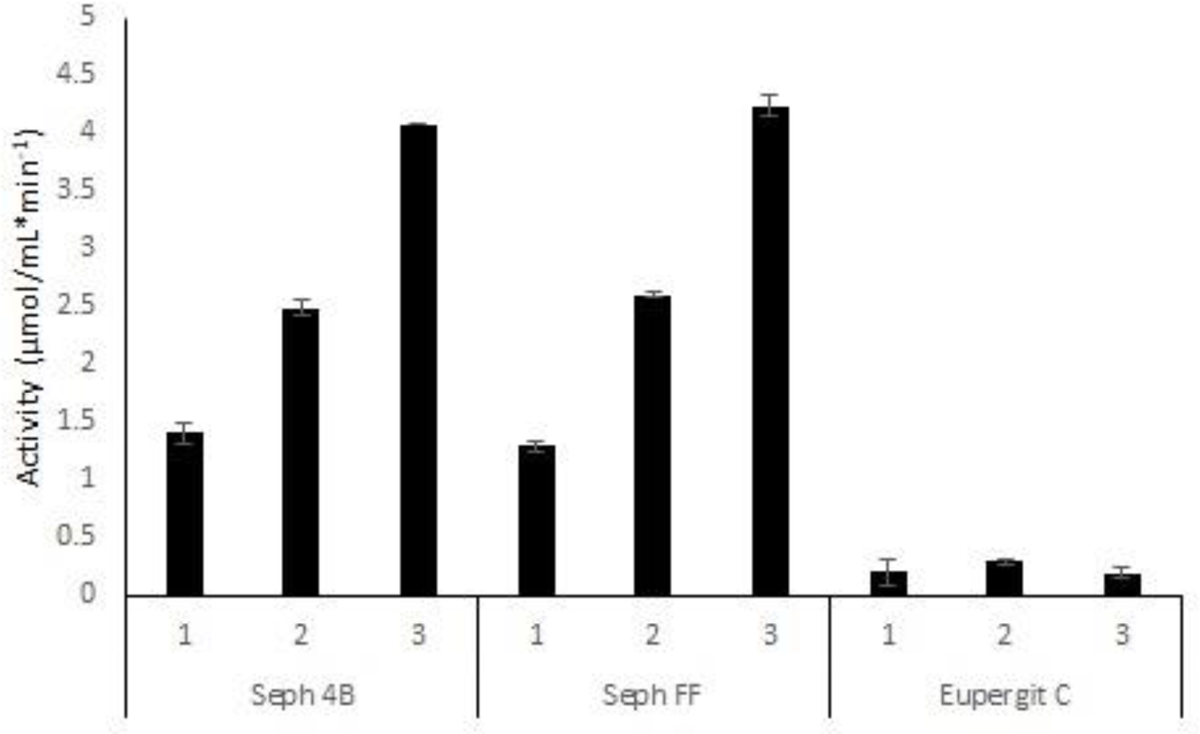
Effect of loaded protein amount on the enzymatic activity of the immobilized enzyme AmyA upon the different supports. Three different protein amounts were used during immobilization: 1, 180 U; 2, 360 U, and 3, 900 U on 50 mg of either Sepharose 4B, Sepharose FF, or Eupergit C. The activity shown corresponds to the initial velocity of reducing sugars released, expressed as the equivalent of glucose when using a fortieth of the respective immobilized enzyme.

### Alcoholytic activity of immobilized Amy A

Unlike hydrolytic reactions, alcoholyis reactions take longer (12 to 24 h) and require higher substrate concentrations (10% starch) to favor the transfer reaction and yield detectable products. Therefore, kinetics could not be followed under our experimental conditions. The butanolysis reactions were performed with 1.6 or 3.9 U of immobilized enzyme at 85 °C for 24 h. At the end of the reaction, butyl glucoside and butyl maltooligosaccharide products were detected by TLC. The spots corresponding to the alcoholysis products before treatment with amyloglucosidase can be observed in the upper part of the TLC plate in the left panel of Figure 2A. Furthermore, by using a glucose standard as reference, the starch hydrolysis products are also visible. The lack of standards of all possible butyl oligosaccharides hindered their quantification; nevertheless, to evaluate the alcoholysis efficiency, the reaction products were hydrolyzed with amyloglucosidase, which cleaves α-1,4 glycosidic bonds from the non-reducing end (22). After digestion with amyloglucosidase, most alcoholysis products were converted to butyl glucoside, and quantified as representing the total alcoholysis efficiency, as shown in the right panel of Figure 2A. Other hydrolysis products such as maltodextrin, were also reduced to glucose. Only a slight difference in the total production of butyl glucoside between the Eupergit C (4.0 mg/mL± 0.2) and Sepharose (6.1 mg/mL± 0.01) was detected (Figure 2B). Even though the hydrolytic activity of immobilized protein increases with the amount of protein loaded on the support, this is not reflected in the final alcoholysis productivity, since this is determined after 24 h, once the reaction reaches equilibrium. In consequence, the initial concentration of the enzyme used for immobilization did not affect the yield of alcoholysis. However, an effect due to the initial enzyme concentration after reusing the support becomes clear, as shown in Figure 3. The concentration obtained in the first cycle was taken as 100 % for each condition, following the production of butyl glucoside over the course of sequential alcoholysis cycles. As a result, the alcoholysis performance of the immobilized AmyA WT on Sepharose FF biocatalyst resulted in a higher yield (29.9% ± 0.15 of butyl glycosides after 6 cycles) compared to the enzyme immobilized on Eupergit C (8.8% ± 1.9 after 4 cycles).

**Figure 2.**
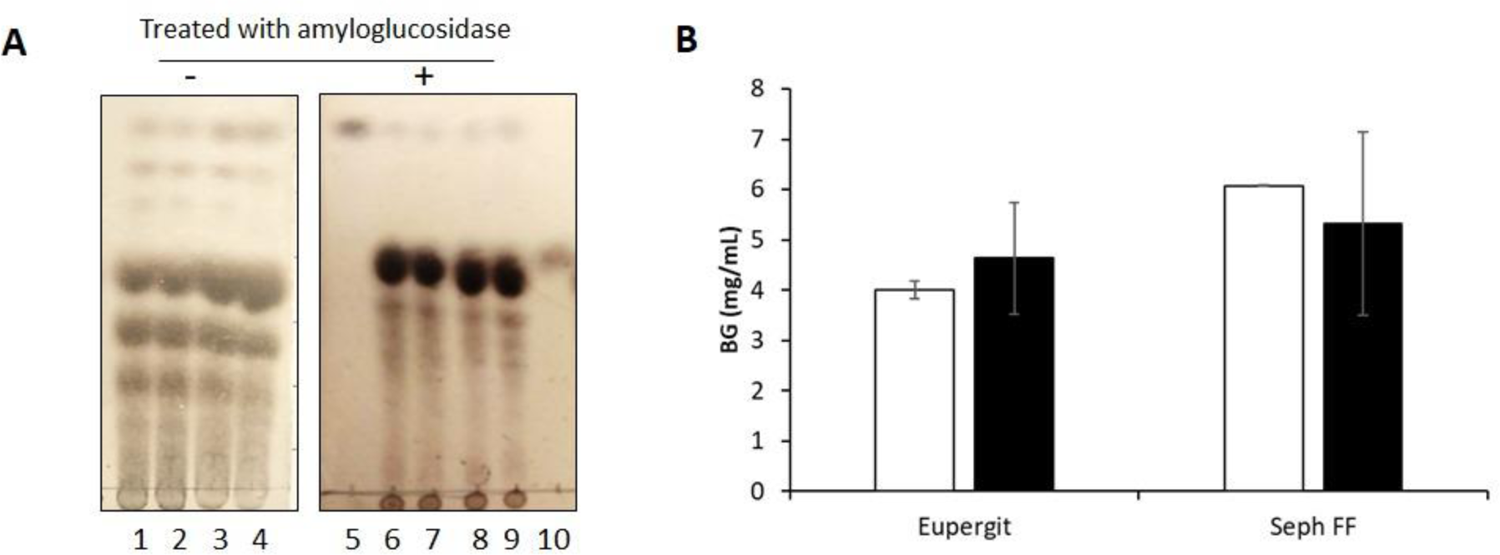
Comparison of the butanolysis reaction products with immobilized AmyA WT. A) TLC of the final products from the butanolytic reaction before (-) and after (+) treatment with glucoamylase (see text for details). Lanes: 1 and 6 = Eupergit C, low-protein concentration; 2 and 7 = Eupergit, high-protein concentration; 3 and 8 = Sepharose FF, low-protein concentration; 4 and 9 = Sepharose FF, high-protein concentration; 5 = butylglucoside standard; 10 = glucose standard. B) Butyl glucoside (BG) title quantified by HPLC after treatment with glucoamylase of the reactions described in A. The white and black bars correspond to low (106 U) and high loaded protein (258 U) in 50 mg of support, respectively.

**Figure 3.**
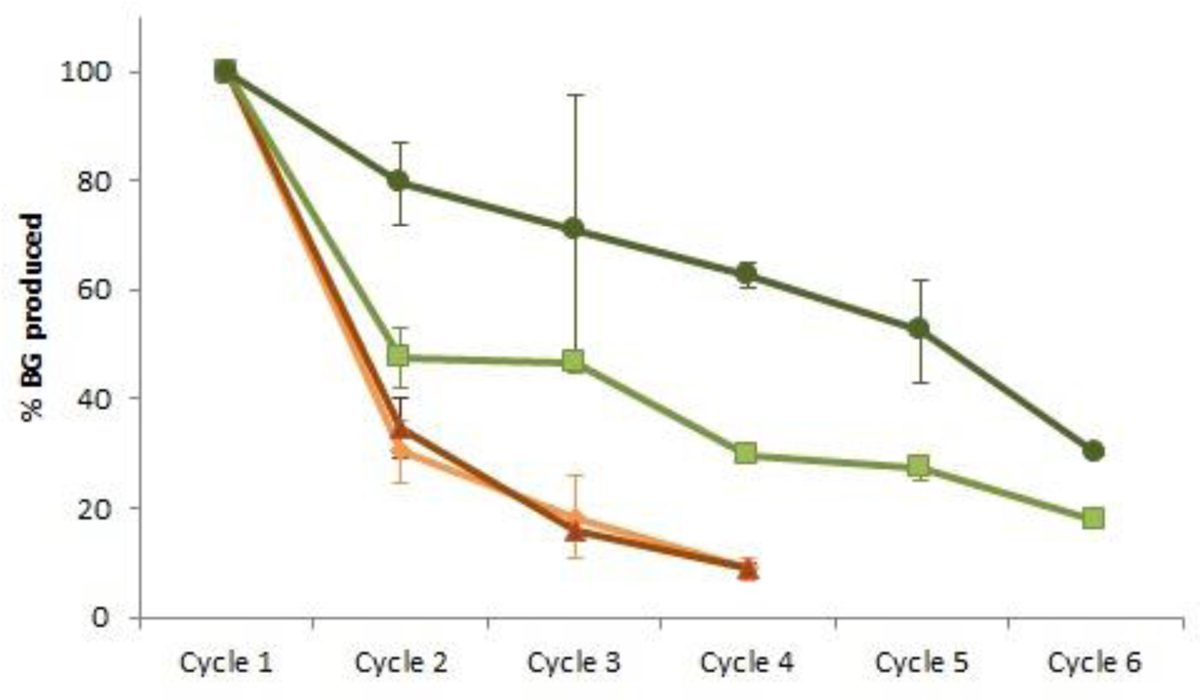
Efficiency of AmyA WT biocatalysts designed with different supports for the butanolysis reaction on different supports. Values for Sepharose FF loaded with 258 U (dark green circles) or 106 U (light green squares) of enzyme are shown. Similarly, Eupergit C loaded with 258 U of enzyme (red triangles) or 106 U (orange diamonds). The percentage is estimated relative to the amount of BG obtained in the first cycle after 24 h reaction for each condition.

### Temperature stability and pH profile of AmyA biocatalyst

When immobilized in Sepharose FF, AmyA successfully carried out the production of butyl glucoside at 85°C (Figures 2 and 3) and showed the best yield per enzyme unit. The Sepharose FF support was then utilized to immobilize the more alcoholytic variant H222Q as well (14). We evaluated and compared the thermal stability of the free and immobilized enzymes at 65 and 85°C after incubation for 6 and 24 h. The residual hydrolytic activity was measured under the optimal reaction conditions, that is 85°C in enzyme buffer at pH 7. The activity was calculated relative to that of the enzyme maintained at room temperature for the same period of time. Interestingly, AmyA WT in the free form showed slightly improved activity after the incubation periods (Figure 4). However, in the case of the immobilized AmyA WT, the increase in activity was almost 50%. Noteworthy, the residual activity of the free WT enzyme is significantly different after 24 h of incubation at 85°C, when compared to the immobilized enzyme. Conversely, the activity of the free H222Q variant was slightly affected by the incubation at high temperatures. However, the immobilized H222Q showed an increased activity comparable to that of the WT enzyme after incubation at high temperatures and retained it even after 24 h of incubation (Figure 4B).

**Figure 4.**
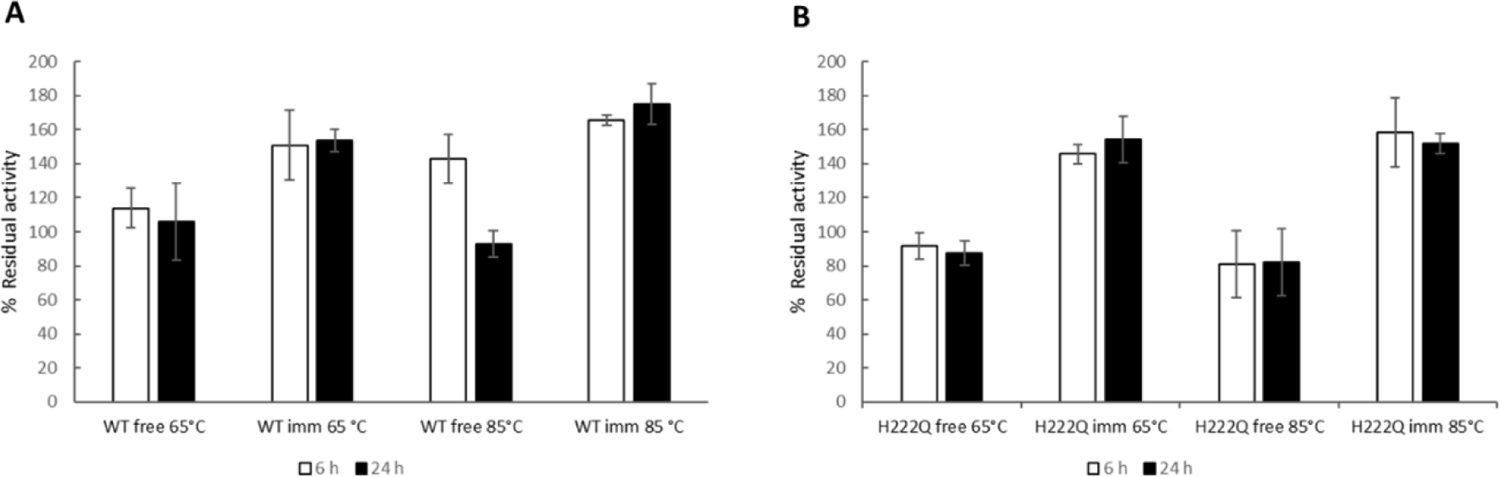
Thermal stability of free and immobilized AmyA WT and the H222Q variant on Sepharose FF. Residual hydrolytic activity values for AmyA WT (A) and H222Q (B). Different incubation times at the shown temperatures were tested (white bars = 6 h incubation; black bars = 24 h incubation). The percentage of residual activity is relative to the activity of the corresponding enzyme maintained after the same incubated period at room temperature.

All activity assays were carried out at pH 7 so far, according to the optimal pH reported for AmyA (23). Nevertheless, we decided to evaluate the effect of different pH values on our immobilized versions of AmyA. For this purpose, we measured the hydrolytic activity in a pH range from 5 to 10.5 while keeping the temperature constant at 85 °C. As a control, we compared the observed activity values with those obtained for the free enzyme at its optimal pH. Figure 5 shows the pH profile for Amy A WT (Figure 5A) and the H222Q variant (Figure 5B). From our experiments, the optimal pH for the free AmyA WT was found to be 7.6. Such value slightly shifted to 7.8 in the immobilized enzyme, and the overall profile showed a minor shift towards higher pH values. The H222Q variant, on the other hand, barely showed a change on the optimal pH from 7.5 to 7.6 upon immobilization, with a rather similar profile between both free and immobilized forms.

**Figure 5.**
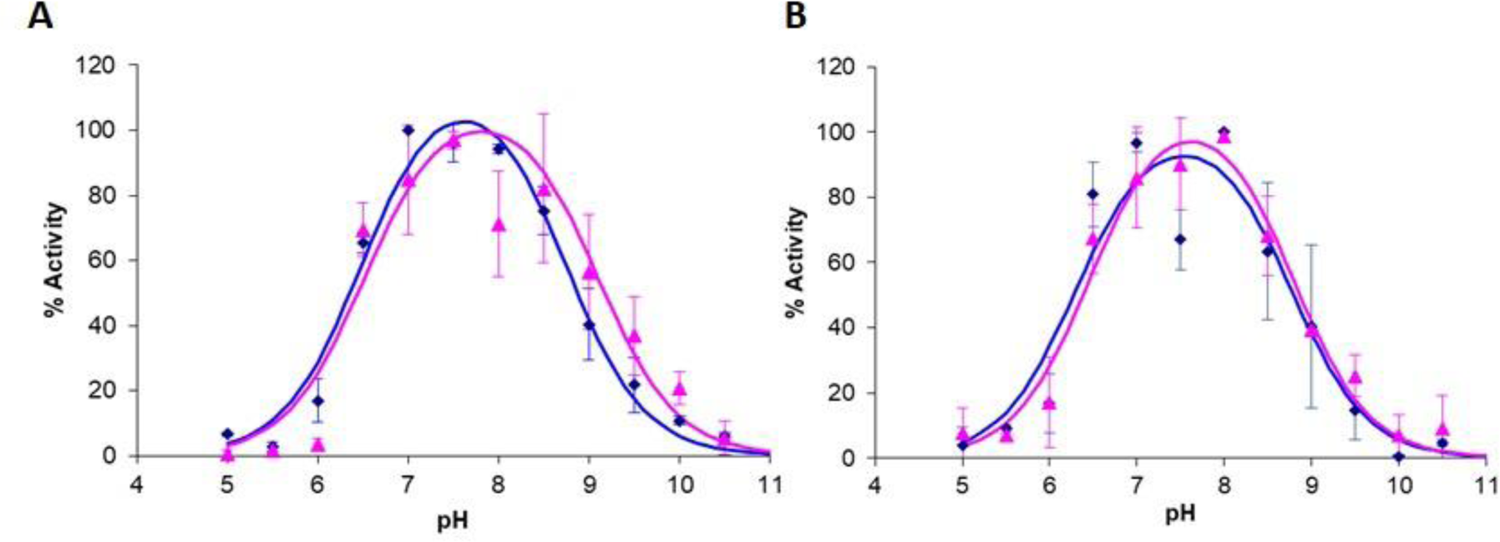
pH profile of free and immobilized AmyA WT and H222Q. Percentage of the hydrolytic activity of AmyA WT (A), and H222Q (B), relative to their free forms at pH 7. Blue lines correspond to the free enzyme, while magenta lines represent the enzyme immobilized in Sepharose FF.

### Synthesis of alkyl glucosides from longer chain fatty alcohols

We investigated the synthesis of long-chain alkyl glucosides with immobilized AmyA by using *n*-hexanol, *n*-heptanol, and *n*-octanol as acceptors. In comparison to the alcoholysis assays with butanol (which represented 10% of the reaction volume), the reactions with *n*-hexanol and *n*-octanol were tested with 5, 10, or 20% V/V of the total reaction volume. The solubility of these alcohols in water is very low, with higher concentrations inducing the separation of phases (aqueous and organic), where the free enzyme migrates towards the aqueous phase. Even though this could favor the hydrolysis reaction, the alcoholytic reaction is rather limited. For this reason, and aiming for experimental reproducibility, we established 5% alcohol in the reaction volume. We had previously observed that the use of polyethylene glycol (PEG) as a co-solvent helped to emulsify all components in the reaction reducing the formation of two phases. Strikingly, in the absence of PEG, no octanolysis products were observed (data not shown). As qualitatively observed by TLC analysis (Figure 6A), the results of the alcoholysis reactions in the presence of PEG were successful in terms of the formation of the expected products, although in smaller amounts compared to the butanolysis (Figure 2A). Although the reactions were incubated for 24 h to reach equilibrium, there was not a clear difference in the product yield when either the free or the immobilized enzymes were used. Nevertheless, the H222Q variant resulted in the highest alkyl glucoside production (lanes 3 and 4 in Figure 6A), similar to previous results with methanol (14). In fact, by increasing the amount of alcohol up to 20% (V/V) in the reaction (with more vigorous stirring), the immobilized H222Q variant produced the corresponding alkyl glucosides even in the absence of PEG (Figure 6B).

**Figure 6.**
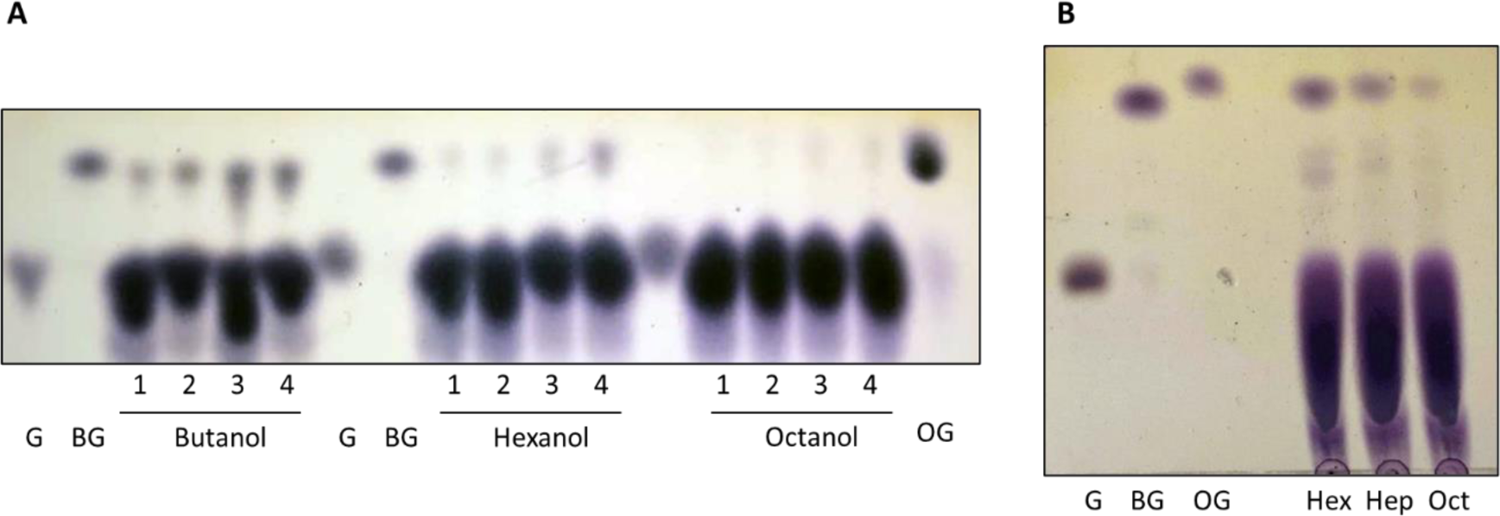
TLC of Alcoholysis reactions carried out by free and immobilized AmyA WT and H222Q. A) Reactions in the presence of PEG: G = glucose standard, BG = butyl glucoside standard; OG = octyl glucoside standard. For all alcohols, lane 1= free AmyA WT; lane 2= AmyA WT immobilized on Sepharose FF; lane 3 = free H222Q variant; lane 4 = H222Q variant immobilized on Sepharose FF. B) Reactions in the absence of PEG utilizing 20 % of the corresponding alcohol with the immobilized H222Q variant. Hex = n-hexanol; Hep = n-heptanol; and Oct = n-octanol.

Octyl glucoside (OG) is a high-value surfactant. Therefore, efforts to increase its yield are worthwhile. To provide an emulsion rather than a two-phase reaction system, we decided to investigate the effect of adding co-solvents to increase the availability of octanol as acceptor in the reaction medium in a similar manner as before. Here, we compared the production of OG in the presence of (1) DMSO, (2) a mixture of DMSO and PEG, or (3) in the presence of other organic alcohols that could serve as co-solvents, such as propanol or butanol. Production of OG was quantitatively estimated by HPLC after treatment with amyloglucosidase to reduce all the octyl glycosides to octyl glucoside. The OG concentration obtained in the absence of co-solvent was defined as 100% for a quantitative comparison with the other conditions (Figure 7). In spite of no significant differences in alcoholysis yield between free and immobilized AmyA (when compared under the same conditions), the addition of co-solvents indeed favored the OG synthesis. In effect, in the presence of PEG up to 7 and 10 % more OG were produced with both free and immobilized AmyA WT, respectively. The presence of butanol and propanol also slightly increased the OG production, particularly when immobilized AmyA WT was used as biocatalyst. Noteworthy, butanol and propanol are both nucleophiles able to act as acceptors, so butyl and propyl glucoside could also be present as products but were not detected in the octyl glucoside HPLC analysis conditions, as they are masked by the presence of DMSO. Surprisingly, the addition of DMSO augmented the OG production by approximately 50 % with both free and immobilized AmyA WT. Finally, as an attempt to evaluate their combined effect, the addition of PEG and DMSO together was not different than the reaction containing only DMSO. Therefore, the use of DMSO as a co-solvent was further investigated for octanolysis reactions with the H222Q variant (Figure 8). Ultimately, there is a clear qualitative difference between the synthesis of OGs produced under these conditions and the reaction without any co-solvent shown in Figure 6. As determined with DMSO, the yield of OG produced with the immobilized variant H222Q is higher than the yield obtained with WT. However, in both cases, they are independent of the enzyme activity units loaded.

**Figure 7.**
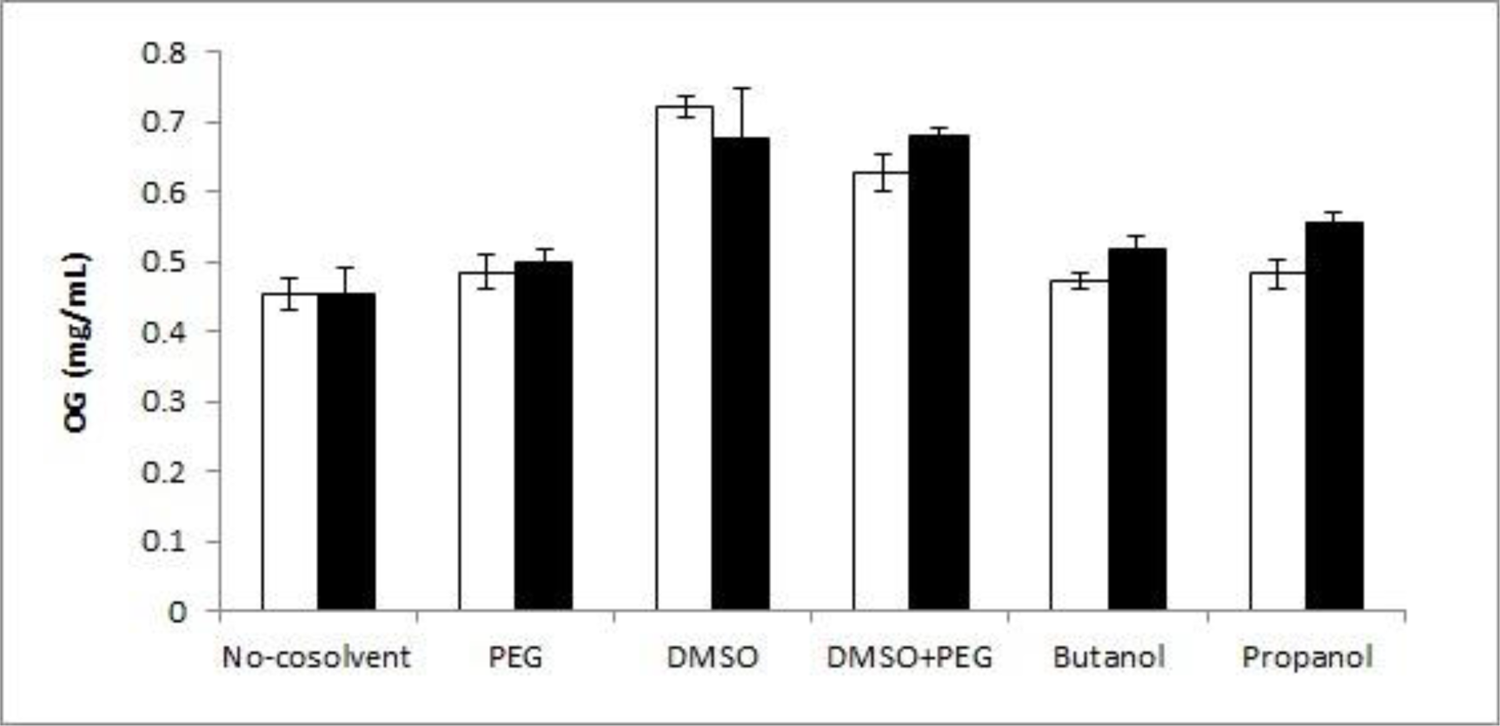
Effect of co-solvents added for the production of octyl glucoside (OG) with immobilized AmyA WT. Alcoholysis reactions with free and immobilized AmyA WT in the presence of 10% octanol and different co-solvents: PEG (5%), DMSO (30%), propanol (5%) or butanol (5%). The concentration of OG was estimated by HPLC after treatment with glucoamylase. White and black bars correspond to the reaction performed with the free and immobilized AmyA, respectively.

**Figure 8.**
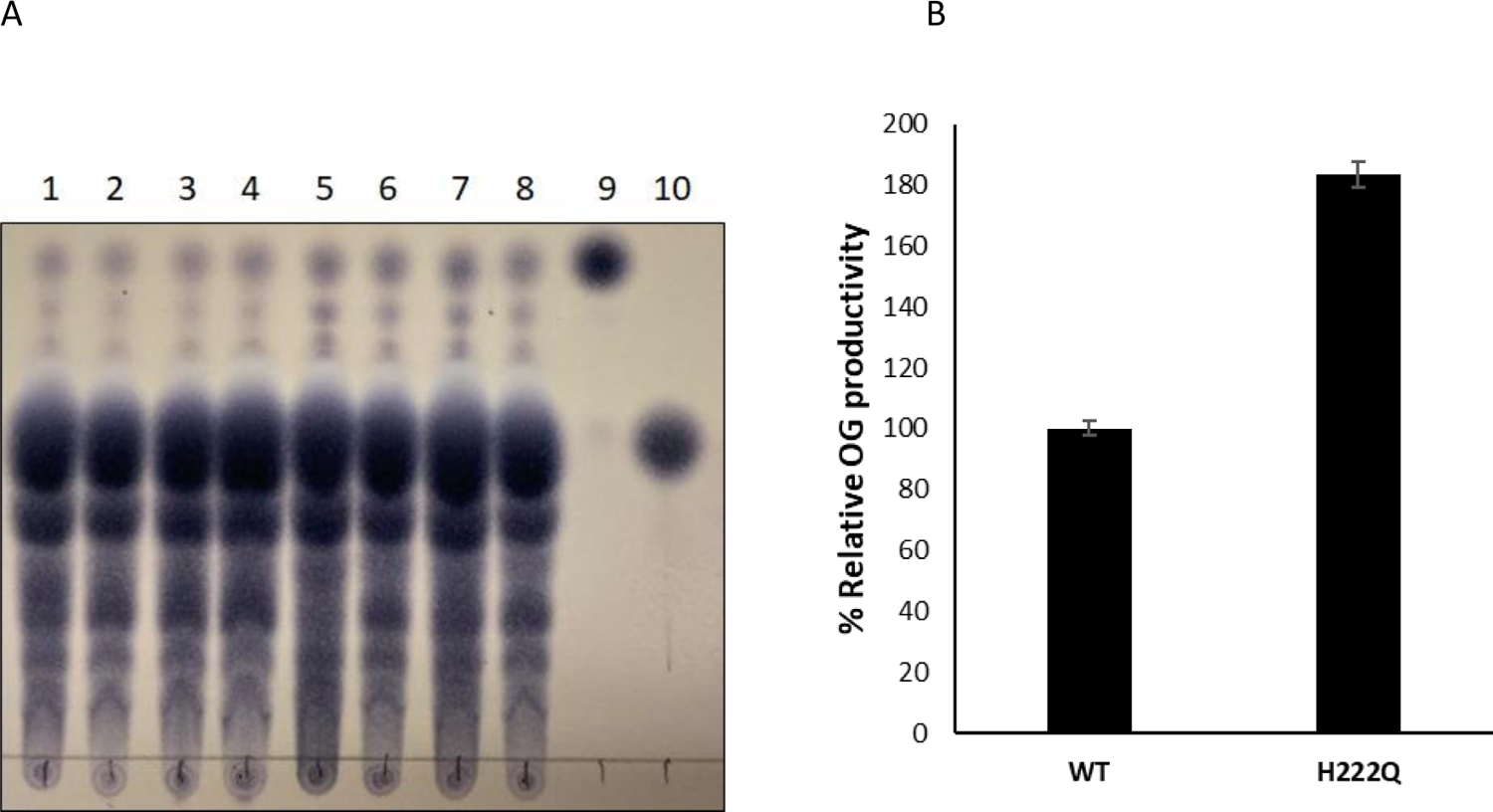
Octanolysis reactions in the presence of DMSO by immobilized AmyA WT and H222Q variant. (A) TLC of the octanolysis reactions in the presence of 20 % octanol and 30 % DMSO. The samples were resolved by TLC before treatment with amyloglucosidase. Lanes 1 and 2 = replicate with 10 µL immobilized AmyA WT; lanes 3 and 4 = replicates with 20 µl immobilized AmyA WT; lanes 5 and 6 = replicates with 10 µL immobilized H222Q; lanes 7 and 8 = replicates with 20 µL immobilized H222Q; lane 9 = OG standard; lane 10= glucose standard. (B) Quantification of octanolysis reaction products after reducing Octyl poly-glucosides to octyl glucoside by amyloglucosidase. The OG production is relative to that of immobilized enzyme under the same reaction conditions.

## Discussion

The implementation of biotechnological processes in the production of APGs through alcoholysis reactions is increasingly desirable from a green chemistry point of view. However, most natural enzymes do not perform at high alcohol concentrations. From this perspective, the use of enzymes obtained from microorganisms that grow under extreme conditions guarantees their high stability, which hopefully can be extrapolated to other adverse conditions beyond high temperatures, such as the presence of alcohols. We have successfully accomplished the production of alkyl glucosides with the thermotolerant α-amylase from *T. maritima* (AmyA). This enzyme has been already proven for alcoholysis reactions withstanding up to 30% methanol (14). In this study, we further explored AmyA increasing its stability through immobilization. Although Eupergit C (an enzyme support) promised to be optimal for enzyme immobilization, given its applications in industry (19), was not the best option, as observed by a significant loss of enzymatic activity upon enzyme immobilization (Table 1 and Figure 1). The use of Sepharose FF, instead, efficiently bound and AmyA with an activity immobilization yield of 96-98%, and an enzymatic efficiency of 29-30%% (Table 1 and Figure 1). Even though immobilization leads to a loss of activity, this is compensated by the possibility of reutilizing the enzyme during several reaction cycles. Strikingly, the immobilized enzyme retained above 50 % of alcoholysis activity after 5 reaction cycles (Figure 3), allowing the production of > 20 mg/mL of BG, in contrast to the previously reported 0.76-0.82 mg/mL titles obtained with Taka-amylase (22, 24). Although the amount of enzyme loaded does not affect the yield of BG during the first reaction cycle, loading higher amounts of protein maintains higher residual activity after several reaction cycles (Figure 3). Furthermore, the enzymes immobilized on Sepharose FF showed a 20 % increment in the production of BG relative to the free enzyme (14) and that of the immobilized on Eupergit C (Figure 2B). This increased title could be explained by the stabilization of the enzyme upon immobilization. The alcoholysis reaction is kinetically controlled, and long reaction times are required to accumulate alkyl glycosides (25). The free enzyme loses activity after 24 h of incubation at 85°C as shown in Figure 4A. This loss of activity can have a negative impact on the yield of alcoholysis products, where the reaction takes longer. On the other hand, the immobilized protein does not show any loss of activity. As a result, a higher amount of alkyl glucosides is obtained after 24 h of reaction.

The H222Q variant, previously engineered to increase the alcoholysis/hydrolysis ratio (14), particularly benefited after immobilization. As shown in Figure 4B, the free form of this variant is less stable than AmyA WT, and loss of activity is observed when the enzyme is incubated at temperatures as low as 65°C for periods as short as 6 hours. Immobilization not only maintained the initial activity (as measured by the hydrolysis of starch) but also increased the activity relative to that of the enzyme maintained at room temperature. This increase in activity upon incubation at high temperatures is indicative of a possible gain in flexibility, leading to an improved accessibility of the enzyme upon immobilization. Thus, incubation of the enzyme at high temperatures allows for the relaxation of regions that may have been restricted while immobilized, resulting in an increase in activity. This is also consistent with the increase in Topt commonly observed in other immobilized proteins (24, 26, 27).

In terms of pH, the immobilization had a larger effect on Amy A WT, with a small broadening towards higher pHs. However, this effect is smaller when compared to other proteins that show a dramatic pH profile change upon immobilization (26, 28).

Strikingly, the combination of co-solvents with longer-chain alcohols increased the alcoholysis production achieved by immobilized AmyA - especially when DMSO was used-producing up to 2.5 mM (0.72 mg/mL) of OG in a single cycle. It has been suggested that DMSO, a denaturant, may contribute by increasing the flexibility of the enzyme molecules that were constrained due to protein-protein interactions in an anhydrous organic medium (29). This is a plausible explanation, consistent with the observation of an increased activity of the immobilized enzymes after incubation at high temperatures. However, it is also possible that DMSO polarity contributes to the solubilization of long-chain alcohols increasing their availability as acceptors in the reaction media.

In contrast, the production of alkyl glucosides by glycosyl-hydrolases has been investigated before with members from families 1 (25, 30, 31) and 3 (32), specifically some β-glucosidases. Nevertheless, *p*-NP glucoside was used in these studies as the glucoside donor and not only represents an expensive substrate, but the yields decreased considerably with longer alcohol chains.

In the present study, the immobilization of the thermophilic AmyA WT and its H222Q variant – with improved transglycosylation activity – conclusively resulted in an increased thermal stability as measured by their hydrolytic activity even after 24 h of incubation at 85°C. In combination with the use of DMSO as co-solvent, a higher production of alkyl glucosides is observed. Of particular importance, our immobilization strategy improved the use of long-chain alcohols such as n-octanol for the biosynthesis of XX mg/mL of octyl glucoside. To our knowledge, this is among the highest titles reported for octyl glucosides. Noticeably, the ability to use an extremely abundant and unexpensive substrate such as starch, positions AmyA as a promising biocatalyst for the production of alkyl glucosides.

## Author Contribution Statement

W.X-V. Conceptualization, Funding acquisition, writing manuscript, experimental methodology, data analysis, F.O.R-A, experimental methodology, editing manuscript A.M-M experimental methodology, data analysis, A.L-M, resources, Writing - Review & Editing, L.O-R, experimental methodology, GSR Conceptualization, Funding acquisition, Writing - Review & Editing

## Acknowledgment

The authors are indebted to Fernando González-Muñoz for assistance in the HPLC determinations, to Ma. Elena Rodríguez Alegría for assistance with Eupergit immobilization, and Dr. Humberto Flores for technical support (Departamento de Ingeniería Celular y Biocatálisis, Instituto de Biotecnología, UNAM), and Juan Manuel Hurtado Ramírez and Roberto Rodríguez, Servando Aguirre and David Castañeda (Unidad de Cómputo Instituto de Biotecnología, UNAM), for their computational support as well as Shirley Ainsworth and Omar Arriaga, for librarian assistance.

## Funding

This research was funded by DGAPA-UNAM through PAPIIT Grant No. IN202619 to W.X.V. and IN226623 to G.S.R.

---

**Figure S1.**
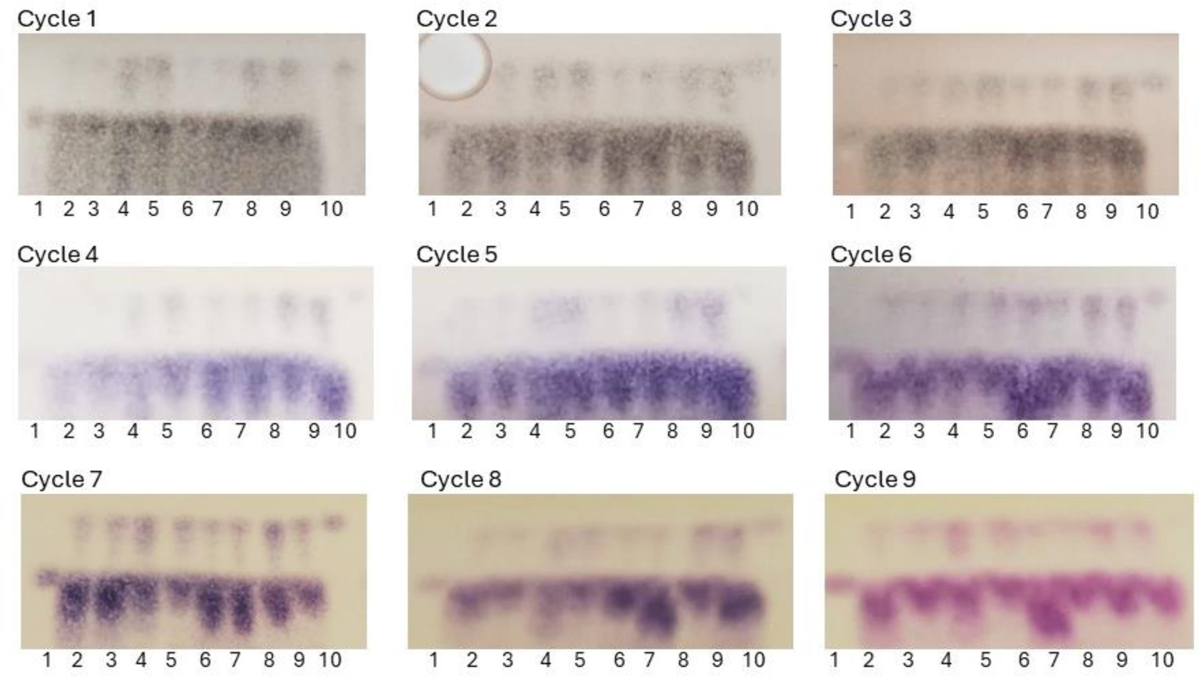
Butanolysis reactions with AmyA WT immobilized in non-crosslinked Sepharose 4B during several cycles. Lane 1= Glucose standard, lane 2 = Free WT, lane 3 = Immobilized WT, lane 4 = Free H222Q, lane 5 = Immobilized H222Q, lane 6 = Free WT, lane 7 = Immobilized WT, lane 8 = Free H222Q, lane 9 = Immobilized H222Q, lane 10 = Butyl glucoside standard. The reactions were incubated at 60°C, lanes 2-5 for 6 h, and lanes 6-9 for 24 h.

